# The Application of Countercurrent Chromatography for the study of Bacteriophages

**DOI:** 10.1101/2021.01.08.425911

**Authors:** Jessica C A Friedersdorff, Colin Bright, David Rooke, Christopher J Creevey, Alison H Kingston-Smith

## Abstract

Bacteriophages (phages) are viruses that target bacteria, with the ability to lyse and kill host bacterial cells. Due to this, they have been of some interest as a therapeutic since their discovery in the early 1900s, but with the recent increase in antibiotic resistance, phages have seen a resurgence in attention. Current methods of isolation and purification of phages can be long and tedious, with caesium chloride concentration gradients the gold standard for purifying a phage fraction. Isolation of novel phages requires centrifugation and ultrafiltration of mixed samples, such as water sources, effluent or faecal samples etc, to prepare phage filtrates for further testing. We propose countercurrent chromatography as a novel and alternative approach to use when studying phages, as a scalable and high-yield method for obtaining phage fractions. However, the full extent of the usefulness and resolution of separation with this technique has not been researched; it requires optimisation and ample testing before this can be revealed. Here we present an initial study to determine survivability of two phages, T4 and ϕX174, using only water as a mobile phase in a Spectrum Series 20 HPCCC. Both phages were found to remain active once eluted from the column. Phages do not fully elute from the column and sodium hydroxide is necessary to flush the column between runs to deactivate remaining phages.

## 2. Introduction

### 2.1. Bacteriophages

Bacteriophages (phages) are viruses that infect only bacterial cells. Once bound to the bacterial surface, the phage genomic material enters the cell and is either incorporated directly into the bacterial host DNA, where it remains for future generations of the host, or the phage genome is replicated and many more virus particles are produced and the cell eventually bursts, releasing the virions to infect other cells. The latter is called the lytic cycle, and because this causes rapid death of the host cell, has become an interest to scientists as a potential therapeutic [1,2].

### 2.2. Current Methods for Isolation and Purification

With a growing interest in phage research, it is important that a variety of methods are tested to build a repertoire of ways to isolate, separate and purify phages to achieve the standard of preparation required, as well as evaluate cost, time, and yield for each method. A hurdle often encountered by researchers to bring a phage preparation to therapeutic fruition is to abate concerns regarding the safety; requiring a high purity with absence of bacterial toxins [3]. In order to achieve this high standard, typical approaches for small scale purification use centrifugation and filtration to remove any whole bacterial cells from lysates, then removal of bacterial debris and toxins using polyethylene glycol (PEG) precipitation and caesium chloride (CsCl) gradient ultracentrifugation. Although the resulting phage preparations are pure, yields are often low and the method long and time consuming, requiring a trained hand [4]. A scalable method for phage purification is the recent implementation of anion-exchange chromatography. It is presented as an easier alternative to CsCl gradients, but only once the elution profile of the phage of interest has been optimised [5,6]. Recovery yield in the purest elution fraction from this technique varied from 55% up to 99.9% depending on the phage of interest, but no measure of purity was given [5].

Yet even before potential therapeutic preparations are purified, phages still need to be isolated and concentrated, and methods to process large volumes are useful, especially for industrial applications or for sources where phages are sparse. Tangential flow filtration is a common technique used to concentrate viruses from large volumes of water samples, removing bacterial cells and debris particles larger than 0.22μm in size [7]. This allows for large volumes with low concentrations of phages to be processed quickly to concentrate the phages into a lower volume for downstream or further processing or testing, including metagenomic sequencing or isolating individual phages against bacterial hosts of interest.

### 2.3. Countercurrent Chromatography and Field-Flow Fractionation

Counter current chromatography (CCC) is the term often applied to any biphasic immiscible liquid-liquid partitioning technique, with each of the phases acting as either mobile or stationary. CCC has successfully been utilised to separate a range of natural and synthetic target compounds, such as plant extracts and medicinal products [8]. The application of CCC for studying phages in any capacity, to our knowledge, has not been reported in any literature previously.

Another separation technique that can be achieved using techniques similar to CCC is field-flow fractionation (FFF). When an external field is applied perpendicular to a column, compounds that are flowing through the column in a mobile phase interact with this external field. Sedimentation FFF (sdFFF) employs centrifugal forces to elicit separation through differential acceleration. sdFFF can effectively separate particles based on size, volume, mass and density with respect to the mobile phase [9]. It has the power and resolution to separate particles from 1nm to ∼50μm in size, and effectively separated nano and microparticles such as dust, silica beads, sand and soil particles, achieved by simply changing the flow rate [10], offering an intriguing opportunity for virus and phage separation. Studies showed that T4D phage retained infectivity after being exposed to the forces of sdFFF [11] and the separation power of sdFFF was shown when phages T4, T7 and the tobacco mosaic virus were separated from T2 [12]. These early studies show promise that phages can be separated based primarily on size.

Hydrodynamic Countercurrent Chromatography (hdCCC) is a separation technique that employs a planetary centrifuge (J-type). The action of the planetary centrifuge exerts a variable g-force, that results in a large number of mixing and settling stages. Using higher speeds and forces (>250 x*g*) allows much higher stationary phase retention and therefore faster separation times, as used by High-Performance Counter current Chromatography (HPCCC).

### 2.4. Aims

The aim of this research was to apply phages to a Spectrum Series 20 HPCCC, with the ultimate aim to determine separation ability of the hdCCC technique for phages. This preliminary work would not be utilising the HPCCC to its full potential, avoiding the use of a biphasic system in this first instance, instead using the forces the CCC exerts through centrifugal forces and the flow rate, much in the same way of sdFFF techniques. Therefore, the primary property that would separate phages is size, and as such to maximise the probability of achieving clear separation, two well studied coliphages with different size and morphology were chosen; ϕX174 is a microvirus with a capsid diameter of 25 nm, spike proteins and no tail [13], which is smaller than the T4 phage, with an 111 nm long and 78 nm wide elongated icosahedral head, and a 18 nm wide and 113 nm long contractile tail [14]. These phages will be used to model phage activity in the CCC process.

## 3. Materials and Methods

Phages and their respective *Escherichia coli* hosts were obtained from the German Collection of Microorganisms and Cell Cultures (DSMZ); T4 (DSM4505) and host *E. coli* B (DSM613), ϕX174 (DSM4497) and host *E. coli* PC 0886 (DSM13127). Bacteria were propagated using Luria Broth (LB, Melford, Ipswich), either as liquid, or solid with added 2 % agar. LB was supplemented with 5 mM MgSO_4_ to increase phage binding [15]. A 100 μl aliquot of overnight host was mixed with 10 μl of phage suspension, added to 3 ml warm 0.8 % LB overlay agar, and poured over pre-set 2 % LB bottom agar. After incubation overnight at 37 °C, 4-5 ml of Fortier Phage Buffer (FB; 20 mM Tris-HCl, 100 nM NaCl, 10 mM MgSO_4_, [16]) was added to the plates, placed on an angle shaker for >2 hours, before collection into a microcentrifuge tube. These were then spun in a centrifuge at 5000 x *g* for 10 minutes and the supernatant filtered through a 0.45 μm low-protein binding PES syringe filter.

The first runs on the HPCCC machine (Spectrum Series 20; Dynamic Extractions, Tredegar) applied 1 ml of phage sample on the semi-preparative column. Using dH_2_O as the mobile phase, two to three column volumes of the mobile phase was used to flush the column and establish equilibrium. The sample was applied through the sample injector port and the run commenced with a flow rate of 6 ml/min in reverse phase and 1600 rpm. Four UV channels were set to 210 nm, 254 nm, 280 nm and 366 nm, and recorded absorbance during elution, whilst fractions were collected every minute (Foxy R2 Fraction Collector, Teledyne Isco, Lincoln). The run was stopped once the peak was formed and plateaued, the column then flushed with mobile phase, then repeated for the other phage. Samples from fractions either side and from the centre of the peak were taken to test for the presence of phage. First a spot test was carried out in duplicate for both phages, spotting 10 μl of the fraction sample on the respective host; which was grown overnight and 100 μl mixed with 3 ml 0.8 % LB agar and poured over bottom agar and left to set. Once fully dry, the plates were incubated at 37 °C overnight. A 5μl loop was used to take samples from the centre of areas of lysis in fractions 9, 12 and 17 for T4 and 19, 22 and 26 for ϕX174, which were resuspended in a small volume of FB, and tested again using a spot test to verify clearing was caused by phage activity. A concentration gradient spot test was performed by serial diluting fractions using a tenfold dilution, and spotting 10 μl of 10^−3^ to 10^−8^ dilutions onto a square plate with set overlays, formed of 400 μl of overnight culture and 10 ml of 0.8 % LB agar.

0.5 % NaOH was added to the column until the output was alkaline, and left for >2 hours, before flushing with water. Once at neutral pH, 0.1 ml T4 phage sample was added to the column, using water as the mobile phase, 3 ml/min flow rate, in reverse phase and at 1600 rpm. The run was stopped after the peak and the chromatogram had plateaued. Samples were taken from the water input, the water output before phage sample was added, then fractions were collected every 2 minutes, as well as two samples from the column flush once the rotation was off. A spot test on square plates was carried out on one fraction every three, up until vial 19, when each subsequent vial was tested, as well as the control samples collected.

## 4. Results

### 4.1. Phages Retain Infectivity After the CCC Process

The first step to utilising the CCC for phage separation is to evaluate the effects the forces of the process have on the biological activity and viability of the phage. Both T4 and ϕX174 phages survived the process and could successfully infect their hosts using a plaque assay. It is difficult to assess yield or loss of phage virions, as knowing the absolute number of phage particles applied is inherently difficult and can only be evaluated using a plaque assay to give an idea of plaque forming units (PFU), comparing the sample before application to the column to the fractions obtained. It is also difficult to determine whether discrepancies were due to virions inactivation or adherence to the column and lost.

#### 4.1.1. T4

T4 successfully retained infectivity after application to the CCC. A peak was detected at after around 22 minutes of recording (Figure 1A), which is the expected time for the elution of one column volume. Vial 13 had the greatest concentration of phages as shown by the concentration gradient spot assay (Figure 1B,1C). Fractions nine to 17 all tested positive for the presence of phage, with an increase and decrease in concentration matching the peak in absorption in the chromatogram (Figure 1A). Not enough fractions were tested to determine where phage elution began and then stopped. The concentration gradient of the stock T4 sample that was applied to the column revealed plaques visible at 10^−8^, which, when accounting for dilution and sampling volumes, equates to a stock phage solution of ∼10^10^ pfu/ml. The areas of lysis were also confirmed to be caused by phage by observing propagation of lysis patterns in subsamples that were swabbed from the initial spot test and tested again.

**Figure 1.**
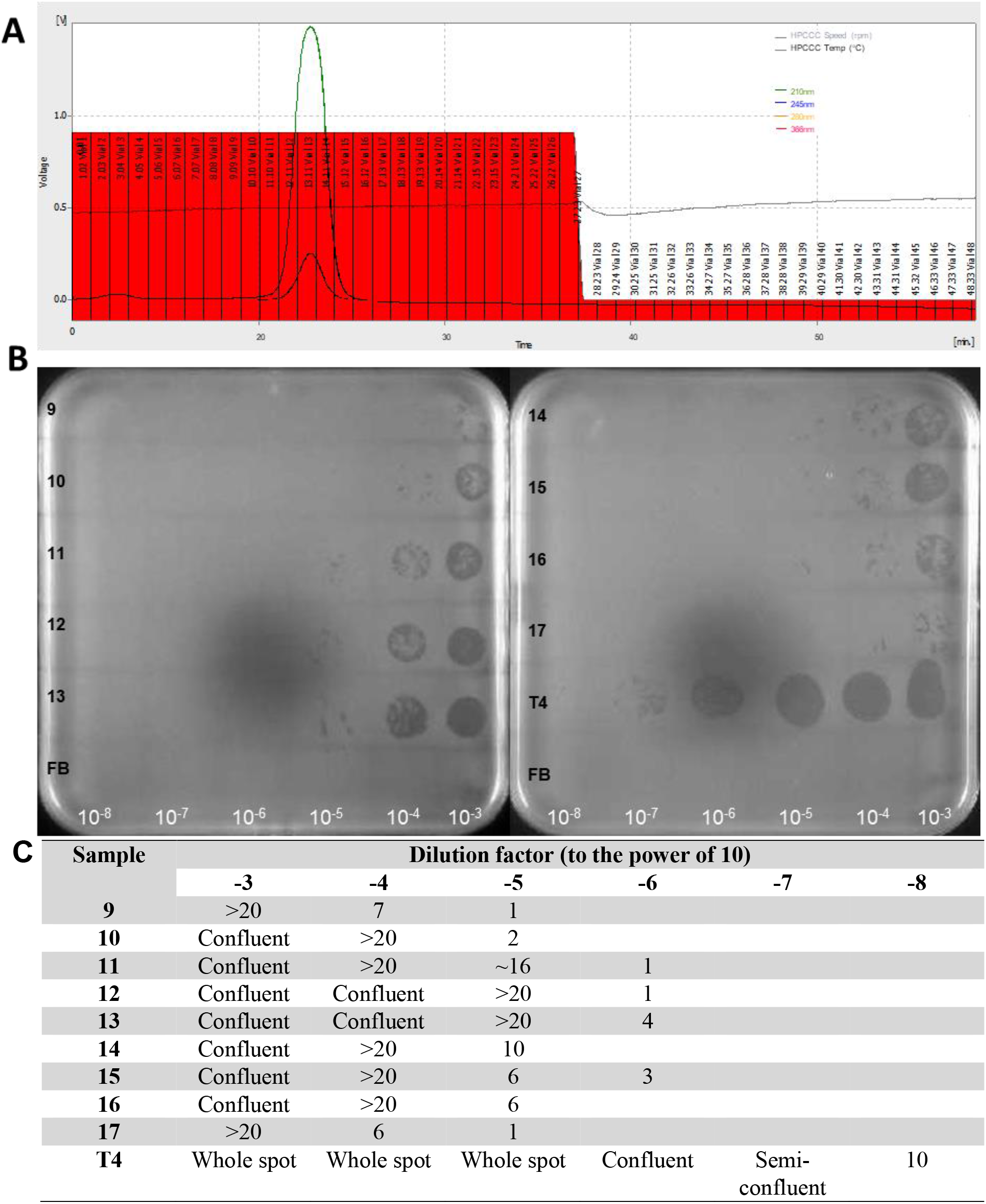
Chromatogram and concentration gradients of the fractions of 1ml T4 sample applied to CCC at 6ml/min. (A) The chromatogram shows the peak at ∼22 minutes and the vial numbers are displayed on the fractions in red bars. (B) Areas of lysis in the form of whole spots, confluent spots or single plaques visible on a bacterial lawn of *E*.*coli* DSM613. Labels of the fraction are in black on the left of the plates and correspond to the vial numbers, and dilution factor in white along the bottom. Fortier Buffer (FB) is applied to the bottom row on both plates as a control. (C) tabulated version of the plates.

#### 4.1.2. ϕX174

Phage ϕX174 also successfully retained infectivity after application to the CCC column. A peak was also seen at around 21 minutes of recording (Figure 2A), which corresponded to one column volume. Vial 22 contained the highest concentration of phages (Figure 2B, 2C), which corresponds to the tail end of the peak, not the centre (Figure 2A). All fractions contained phages, as seen by presence of areas of lysis in a spot test (data not shown) but as with T4, not enough fractions were tested to fully encapsulate the start and end of the elution of the phage sample. No areas of lysis or plaques were seen for fraction number 26 on the dilution gradient, but given the area of lysis present on the initial spot test for this fraction (data not shown), this would suggest a concentration of phages in the fraction of less than 10^5^ PFU/ml. The concentration gradient revealed that the initial concentration of the phage solution was ∼10^10^, when calculating for dilutions. As with the T4 samples, the areas of lysis were indeed caused by phage, shown by observing areas of lysis in samples propagated from positive spots.

**Figure 2.**
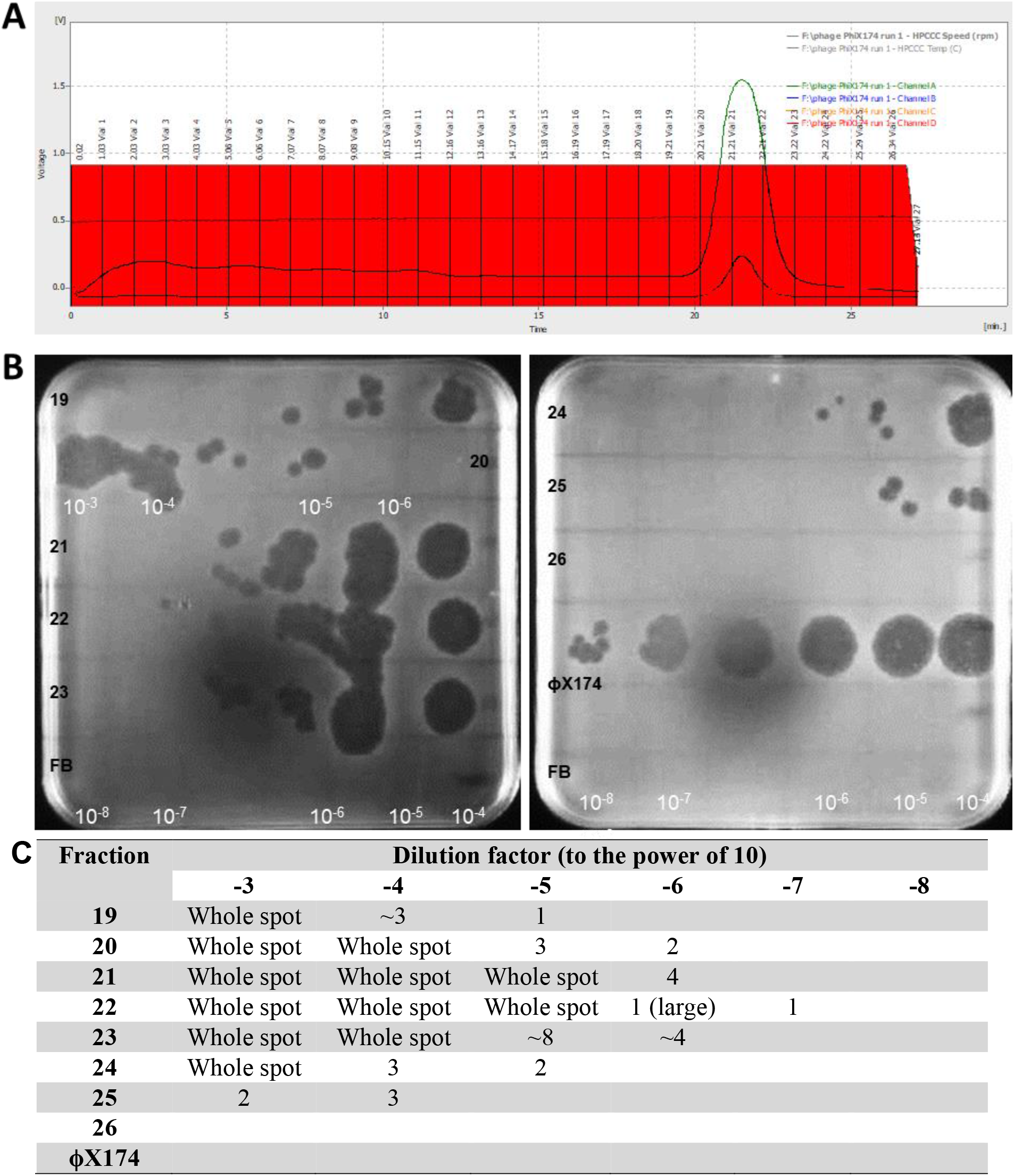
Chromatogram and concentration gradients of the fractions of 1ml ϕX174 sample applied to CCC in reverse phase at 6ml/min. (A) Chromatogram showing the fractions in red with the vial numbers displayed. (B) Areas of lysis in the form of whole spots, confluent spots or single plaques visible on a bacterial lawn of *E*.*coli* DSM13127. Labels of the fraction are in black on the left of the plates and correspond with the vial numbers, and dilution factor in white along the bottom. Sample 20 was spotted incorrectly (human error) and is labelled to correct for this. Fortier Buffer (FB) is applied to the bottom row on both plates as a control. (C) tabulated version of the plates.

### 4.2. Phages that remain in the column are inactivated sodium hydroxide

Additional attempts at loading phage samples onto the column were foiled by the presence of phages from previous runs, revealing lysis in all fractions (data not shown). Leaving the column in a solution of 0.5 % sodium hydroxide (NaOH) for ∼2 hours at the end of a previous run deactivated remnant phages. The NaOH was flushed from the column with the mobile phase before the new phage sample was added. With a flow rate of 3 ml/min, a peak was seen at around 43 minutes, as expected, and no lysis was seen in the spot tests on samples taken from the mobile phase after flushing but before a subsequent phage sample was applied, or in the early fractions before one column volume had eluted (Figure 3). There is no detectable phage activity in the mobile phase before it enters the column, or after it has left the column, which was treated with NaOH. No phages were detected until fraction number 16, where >30 plaques were counted. From vial 19 through to vial 25, the spots showed complete lysis, with confluent lysis visible in vials 26 and 27. A sample of the mobile phase from the flush once the procedure was stopped revealed still a >20 plaques visible. This was reduced to <10 plaques after ∼100 ml of mobile phase was flushed (Figure 3)Figure 3.

**Figure 3.**
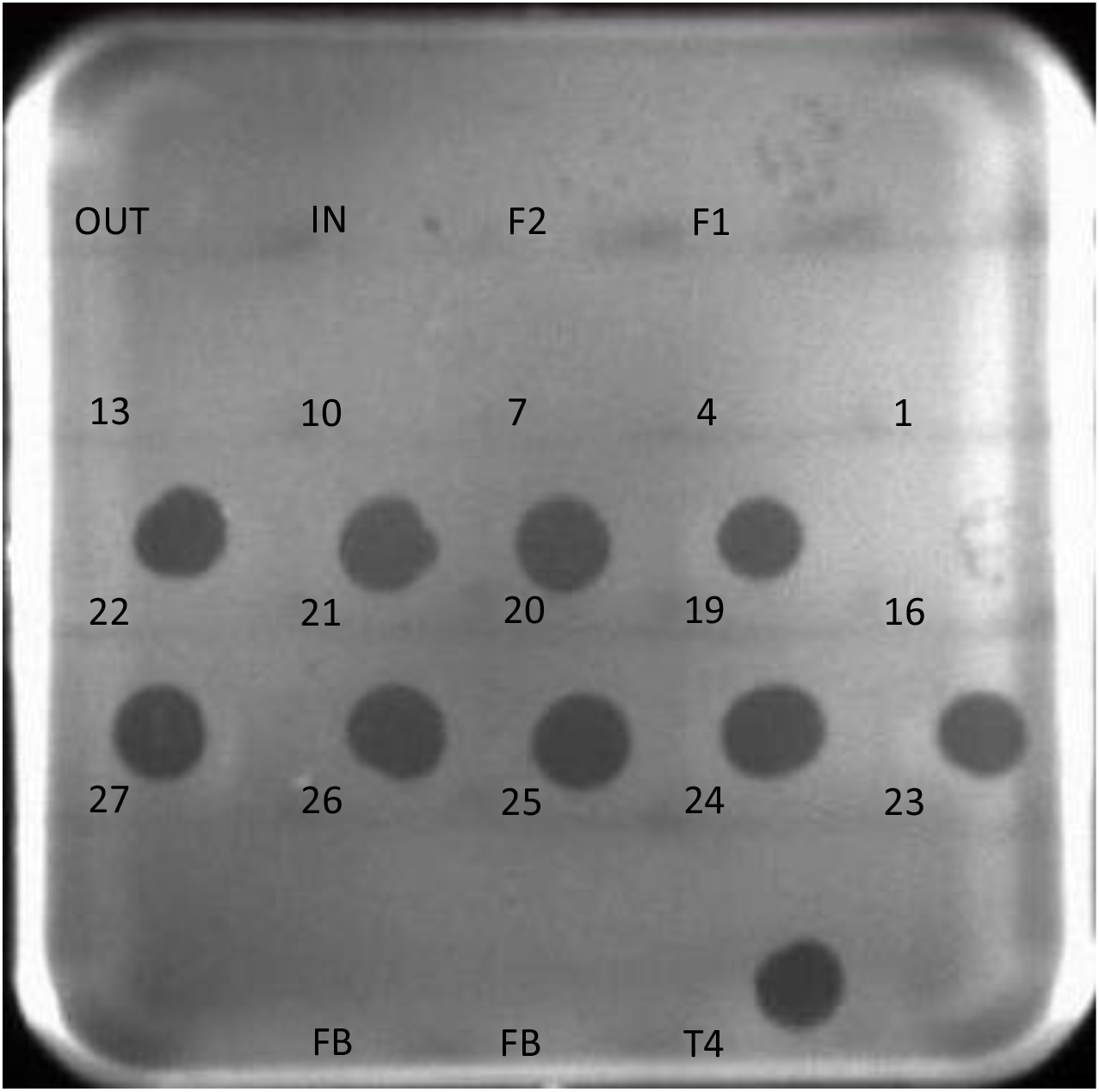
Spot test of fractions of 0.1ml T4 at 3ml/min. The numbers correspond to 10μl of that fraction spotted onto the bacterial host *E. coli* DSM613. IN – a sample of the mobile phase (water) entering the column, OUT – a sample of the mobile phase from the column after NaOH treatment. F1 – column flushed after the revolutions were stopped. F2 – same as F1, but after ∼100ml had been flushed. FB – Fortier Buffer. T4 – the initial T4 sample applied to the column.

## 5. Discussion

Given the proteinaceous nature of the common virus structure, particularly phages, it stands to reason that implementing protein chromatography methods to separate or isolate whole phages directly should be possible. Despite the size difference and more complicated morphology of phages compared to proteins, it is expected that phages would act like proteins in chromatography systems [17]. Therefore, to study the application of field-flow fractionation in a countercurrent chromatography setup is a worthy and novel endeavour.

As with the sdFFF experiments conducted previously, it is important to first establish whether the phages would survive the process [11]. Ideally the phages should remain not only intact but also viable and biologically active, and this activity may be affected by the high shear forces exerted during the process. This was found not to be the case as both T4 and ϕX174 phages could be applied to the column with no ill-effects. Not only this, but the CCC forces did have an effect on the elution profile of the phage samples. Instead of eluting simultaneously (as is injected), there is an increase then decrease in phage concentration which roughly matches the peak seen in the UV detection. With these settings, there was little to no retention of the phages on the column, as phages were detected after one column volume.

Although yield could not be accurately determined using these methods of detection, when a concentration gradient of each of the initial phage samples were spotted onto the respective host and compared to the fractions, there is no significant loss of phages in the resulting fractions. Had a portion of the phages become deactivated or otherwise lost in the column, then it is expected that the concentration of phages in the fractions would be much lower than they are. This is an advantage of the CCC process, since the mobile phase can be fully eluted, and all sample be recovered [18]. Some phages did remain in the column, as shown by plaques formed in the flush collections. This could be evidence of some retention or that there is an interaction between phage particle and PFTE tubing. This will negatively effect yield and is a hurdle to overcome in the future to increase phage recovery.

One of the primary difficulties that this technique will need to overcome is finding a suitable in-line detector that can suitably detect phage particles as they elute. Currently the use of UV and different wavelength channels do not pick up the phages themselves, as the fractions tested corresponded to the peak in the chromatogram but did not encapsulate the start and end of phage elution, and the highest concentration of ϕX174 did not match the fractions corresponding to the peak. Whilst the use of UV detection is also implemented and recommended in the anion-exchange chromatography method for purifying phages [6], there were difficulties in detecting T4, T7 and tobacco mosaic virus elutions from sdFFF using UV detectors, because peaks from the virus particles could not easily be distinguished from background noise [12]. For the current study, to confirm the presence and activity of phages, fractions were spotted onto a lawn of the corresponding *E. coli* host. This method requires more preparation and time, with results available in >6 hours. As long as one active phage is present in 10 μl, then this should be detectable, and therefore theoretically the average lowest level of detection for the spot test is >10^2^ PFU/ml.

Because the method of detection implemented employs the lytic action of the phage to visualise activity on a bacterial host lawn, removing remnant phage particles from previous runs is imperative to avoid false positives. Flushing the column with water or 50 % methanol between runs is not adequate, as shown by the presence of phages in fractions earlier than the solvent front (data not shown). Instead, at minimum, a bolus of 0.5 % sodium hydroxide should be used to flush the column in between runs to ensure remaining phages are inactivated. SdFFF has been often implemented in the separation of biological cells, including human and bacterial [19]. Not only do optimisations need to be made to ensure cell viability, but also high recovery and elution of otherwise sterile fractions. “Channel poisoning” can occur when there are interactions between the particles and the column surfaces, which often arise as the cell suspensions are not pure, similar to the phage samples in this study. This leads to a drop in yield, uncharacteristic elution profiles and reduction of cell viability [19].

## 6. Conclusions and Future Work

In this study, we have shown that the phages T4 and ϕX174 can be applied to a HPCCC column using water as a mobile phase only and remain viable and active after elution. Some phages remained in the column after flushing with mobile phase, making the application of 0.5 % sodium hydroxide necessary to deactivate residual phage particles. Plaque assays proved to be the best method for quantifying phages in fractions, whilst in-line UV detection is not suitable for monitoring phage particle elution. This research paves the way for the utilisation of hdCCC in phage handling, offering promise for separation and purification. Although this research has begun to develop this foundation, further studies are required to determine the resolution of separation this method can achieve by changing the settings and implementing biphasic liquids, such as a polymer-salt system or micelles in an aqueous two phase system, which has been successfully implemented on the filamentous phage M13 [20]. A suitable in-line detector is also required to streamline the process.

## 7. Author contributions

JCAF and CB carried out the laboratory work with advice from DR. The project was conceived by DR, CJC and AKH. All authors contributed to manuscript writing and editing.

## 10. Ethics

There were no ethical considerations needed for this research.

